# Remodeling Activity of ChAHP Restricts Transcription Factor Access to Chromatin

**DOI:** 10.1101/2025.07.03.662926

**Authors:** Josip Ahel, Fabio Mohn, Michaela Schwaiger, Lucas Kaaij, Jennifer Steiner, Eliza Pandini Figueiredo Moreno, Daniel Hess, Marc Bühler

## Abstract

Transcription in eukaryotes is regulated by chromatin-based mechanisms that control nucleosome occupancy, chromatin modifications, and transcription factor binding. We have previously shown that the transcription factor ADNP forms the ChAHP complex with the chromatin remodeler CHD4 and HP1 proteins, acting as a site-specific regulator of transcription and antagonist of CTCF binding. However, the molecular basis of these functions remained unclear. Here, we demonstrate that the CHD4 subunit is essential to antagonize CTCF and silence transcription of transposons, while HP1 proteins are dispensable. Although the remodeling activity of CHD4 is not required for ChAHP chromatin association, it is critical for both transposon repression and CTCF antagonism. Our findings support a model wherein ADNP recruits chromatin remodeling activity in a sequence-specific manner, enabling transcriptional control and local modulation of chromatin architecture.

## INTRODUCTION

Eukaryotic transcription is controlled through the coordinated action of multiple chromatin regulatory mechanisms, including chromatin modifications and chromatin remodeling^1–8^. One principle underlying this regulation is the inability of many transcription factors (TFs) to associate directly with nucleosome-bound DNA^9–12^. Thus, by positioning nucleosomes at key regulatory sequences, cells can regulate transcription in a time- and locus-specific manner. Biochemically, this is achieved by chromatin remodelers which utilize ATP hydrolysis to induce DNA repositioning, nucleosome eviction, or nucleosome incorporation. For example, the NuRD complex deploys chromatin remodeling activity of CHD4 alongside deacetylase activities of HDAC1/2 to facilitate transcriptional silencing^13,14^. In a related yet mechanistically distinct manner, the HIRA and ATRX-DAXX complexes place nucleosomes containing the histone variant H3.3 into chromatin to regulate replication-independent chromatin maintenance^15–17^. Chromatin remodelers themselves normally lack strong sequence specificity, and many are thought to be targeted to specific loci through interactions with TFs. For instance, SNF2H is thought to collaborate with CTCF, inducing an increase in chromatin accessibility^18,19^. Additionally, the remodeler likely liberates CTCF binding sequence motifs from the nucleosome, facilitating CTCF-chromatin binding and resulting in regular nucleosome positioning around these sites. Thus, the functional association of the TF and remodeler is required both for TF recruitment, and for downstream changes in chromatin properties. An equivalent relationship was described for the chromatin remodeler BRG1 and the TF REST^18^. However, in both these and most other cases, the chromatin remodelers are only transiently, weakly, or indirectly associated with the TFs^20–23^.

There are two prominent outliers of this labile recruitment principle: the INO80 and ChAHP complexes. INO80 incorporates the TF YY1, and this interaction is important for sequence-specific targeting of INO80 to gene promoters, resulting in transcriptional activaton^24,25^. Reciprocally, YY1 binding to chromatin is thought to be dependent on INO80^24,26^, mirroring the interdependencies seen with CTCF and REST. ChAHP is composed of the zinc finger TF ADNP, CHD4 and HP1β or HP1γ proteins, which assemble into a stable complex that can be reconstituted *in vitro*^27^. Disruption of ChAHP through deletion of the central ADNP subunit is embryonically lethal, and heterozygous truncations of ADNP result in a severe developmental disorder in humans^28,29^. The most prominent genomic targets of ChAHP are retrotransposons from the SINE family, bound in a sequence-specific fashion^27,30,31^. ChAHP additionally binds heterochromatin, including transposons from the LTR and LINE families, mediated by the association of the HP1 subunit with H3K9 trimethylated nucleosomes^31^. At the molecular level, loss of ADNP, and consequently the ChAHP complex, leads to upregulation of RNA pol III-transcribed SINE B2 elements in mouse embryonic stem cells (mESCs), accompanied by increased chromatin accessibility^27,30^. In addition, many murine SINE B2 elements carry a CTCF motif, and loss of ChAHP results in increased CTCF association with these elements alongside a rearrangement of topologically associated domains^30^. Despite its biological importance and clear link to chromatin biology, the mechanism of action of the ChAHP complex remains unknown.

Several mechanisms have been proposed for ChAHP-mediated genome regulation, including heterochromatin seeding via HP1^32^, competition for chromatin binding with CTCF and other TFs^30^, association with additional remodeler modules such as BRG1^33^, or targeted chromatin remodeling by CHD4^31^. However, none of these models have so far been supported by direct evidence. By generating endogenously edited ADNP mutant mESCs to decouple specific subunits from the complex, here we show that ChAHP-associated CHD4 is necessary for SINE B2 repression, while HP1s are not. We further demonstrate that both SINE B2 repression and CTCF antagonism are specifically effectuated by chromatin remodeling activity of CHD4 within ChAHP. We propose the stable association of ADNP with CHD4 guides ATP-dependent chromatin remodeling activity to locally regulate chromatin.

## RESULTS

### The extreme N-terminus of ADNP interacts with CHD4 to form the ChAHP complex

To test which ChAHP subunits are required to silence SINEs in mESCs, we sought to introduce mutations into the endogenous *Adnp* gene that specifically decouple either HP1 proteins or CHD4 from ADNP. While the interaction with HP1 can be disrupted by introducing point mutations into a conserved PxVxL motif in ADNP, a suitable small mutation abrogating CHD4 binding has not yet been demonstrated (Figure 1A)^27,31,34,35^. To identify such mutations, we used AlphaFold^36^ to predict the structure of the ChAHP complex and infer interaction surfaces between ADNP and CHD4. This revealed that an N-terminal region of ADNP interfaces with a C-terminal region of CHD4, consistent with previously published low resolution interaction mapping (Figure S1A)^27,35^. To validate this prediction, we transiently transfected a series of ADNP truncation constructs into mESCs and probed their interaction with ChAHP subunits by co-IP (Figure S1B). All tested constructs where ADNP N-terminal residues were absent showed strongly reduced CHD4 binding, while HP1γ binding scaled with expression level of the constructs. Reciprocally, all constructs encoding N-terminal fragments of ADNP, including the smallest tested construct (amino acids 1-43), robustly interacted with CHD4. We conclude that the highly conserved first ∼40 amino acids of ADNP are both necessary and sufficient to bind CHD4. To facilitate genome editing and keep the perturbation as precise as possible, we sought to identify a narrower region necessary for binding. A detailed look into the structure prediction revealed that the first 8 amino acids of ADNP form extensive polar sidechain-backbone interactions with CHD4, and one close hydrophobic contact (Figure 1B). We tested the contribution of these residues to CHD4 binding by performing a co-IP with granular truncations of the N terminus (Figure 1C). This revealed that removal of 7 amino acids after the start codon of ADNP results in a comparable decrease in CHD4 binding as the more extensive truncations. Thus, the extreme N-terminus of ADNP is necessary for the ADNP-CHD4 interaction.

**Figure 1.**
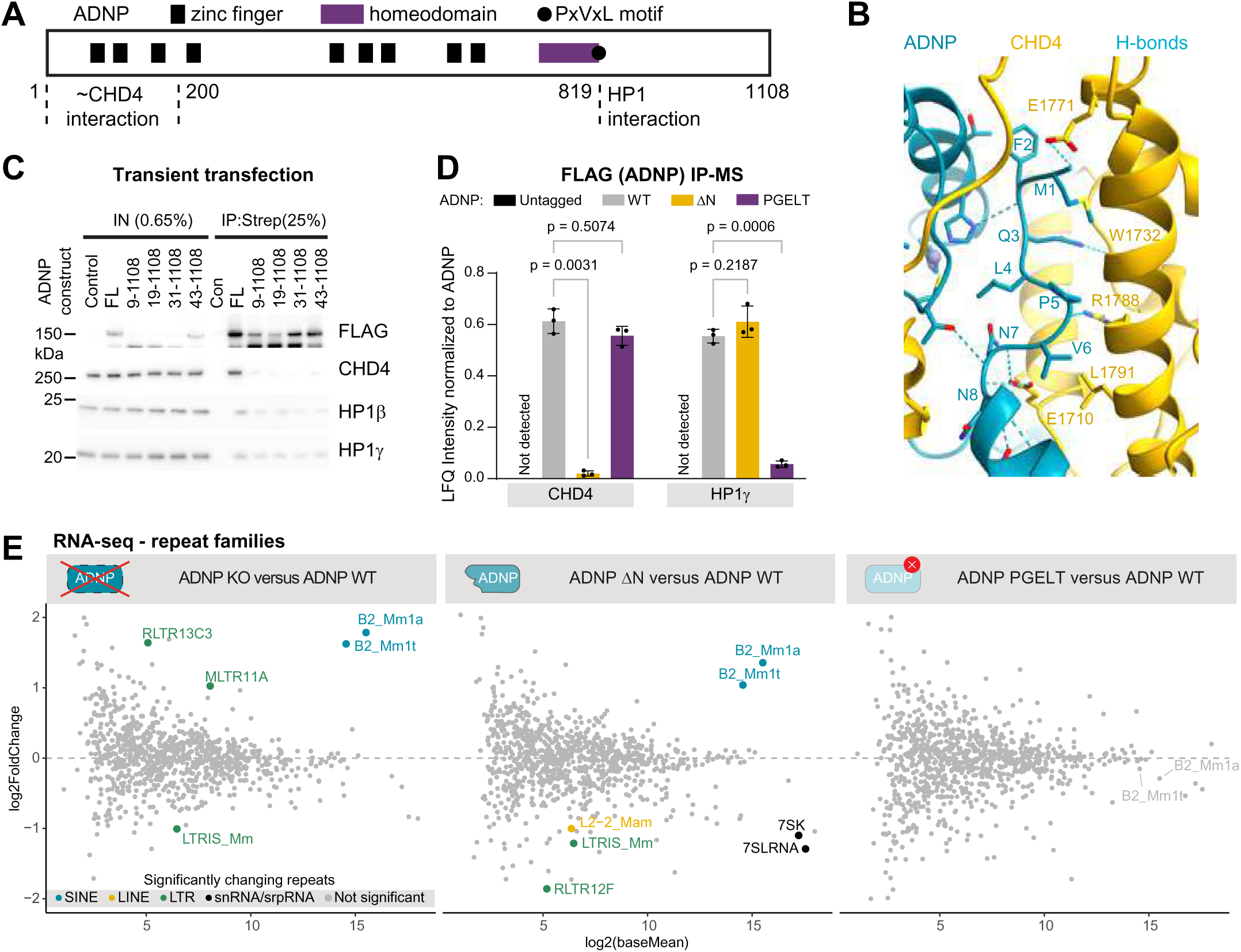
Decoupling CHD4 from ChAHP results in SINE B2 derepression. **(A)** Protein sequence schematic of *M. musculus* ADNP, with key domains highlighted. CHD4 was shown to interact with ADNP with residues corresponding to the first ∼200 amino acids on the ADNP N-terminus. **(B)** Predicted structure of the interface between CHD4 and the extreme N-terminus of ADNP. **(C)** mESCs were transiently transfected with plasmids encoding 3xFLAG-Avi tagged fragments of ADNP as indicated, subjected to co-IP with Streptavidin beads, and analyzed by western blotting. FL = full length. **(D)** Label-free quantification of CHD4 and HP1γ proteins in ADNP IPs, normalized to ADNP intensity (mean ± SD, n = 3). Significance was determined by 2-way ANOVA followed by Dunnet’s multiple comparison test. **(E)** Differential expression analysis for repeat families between ADNP mutants and the corresponding WT control (n=2 for PGELT mutant, n=3 for others). Significant hits are highlighted with colors (FDR < 0.05, |log2FoldChange| > 1). No repeat family is significantly differentially expressed in the PGELT mutant.

We introduced this truncation into mESCs encoding ADNP-Avi-3xFLAG and assessed its effects on ChAHP integrity (Figure 4D, S1C-E). This endogenous intervention fully agreed with the results obtained by transient transfection (Figure S1D). Importantly, the minimal N terminal truncation (hereafter referred to as ADNPΔN) only affected incorporation of CHD4, but not HP1 into the ChAHP complex (Figure 4D). Similarly, auxiliary interactors of the ChAHP complex, such as ZMYM2/3, ATRX or TFIIIC subunits were not affected (Figure S1E). Vice versa, mutating the PxVxL motif to PGELT in the endogenous *Adnp* gene affected HP1 but not CHD4 incorporation. Consistent with a loss of association with heterochromatin, the heterochromatin-associated remodeler DAXX did not co-purify with ADNP-PGELT (Figure S1E). Thus, we were able to generate a panel of genome-edited mESC lines wherein either CHD4 or HP1 proteins were specifically excluded from the ChAHP complex.

### Removal of CHD4 from ChAHP results in SINE B2 activation

We used these cell lines to assess the contributions each subunit makes to the regulation of transcription by RNA-seq and compared them to the effects of complete loss of ADNP (Figure 1E). Consistent with previously published data, SINE B2 retrotransposons were significantly upregulated in ADNP KO cells^30^. Cells harboring the ΔN truncation of ADNP similarly showed a significant increase in SINE B2 RNA level, specifically the less divergent Mm1a and Mm1t subfamilies, while expression of other transposon families was not significantly affected. By contrast, decoupling HP1 from ChAHP induced no significant effects on transposon expression. At the gene expression level, CHD4 decoupling resulted in similar effects as knocking out the *Adnp* gene, and the magnitude of these changes strongly correlated (Figure S1F-G). Conversely, barely any differential gene expression was identified upon HP1 decoupling, and the correlation with effects brought on by *Adnp* KO or ΔN truncation was poor (Figure S1F-G). Together, these data indicate that within ChAHP, CHD4 is required to repress SINE B2s and maintain appropriate gene expression, while HP1 proteins are not.

### ADNP associates with chromatin without CHD4 but inefficiently antagonizes CTCF

Several transcription factors, such as CTCF and REST, rely on auxiliary factors to support their recruitment to chromatin^18^. To test whether ADNP uses CHD4 or HP1 to gain access to its cognate binding sites, we performed ChIP-seq with wild-type ADNP, and both subunit-decoupled mutants (Figure 2A-C, Figure S2A-B). ADNPΔN exhibited increased chromatin occupancy compared to WT, most strongly at ChAHP binding sites overlapping SINE B2 elements, and less so at other ChAHP-bound transposons (Figure 2A-B). Importantly, the capacity of ADNPΔN to bind regions that do not overlap transposons was unaffected. Consistent with previously described effects, decoupling HP1 proteins from ChAHP resulted in an overall increase of ADNP ChIP signal at loci overlapping euchromatin, including SINEs and regions that do not overlap with transposon sequences^31^. In contrast, this increase was not evident at heterochromatic retrotransposons, such as LINEs and LTRs. Since peak calling does not reliably capture broadly distributed heterochromatin-associated ADNP, we supplemented these analyses with a peak-agnostic quantification over transposable elements globally (Figure S2B). Herein, a reduction of heterochromatin occupancy is readily observed upon HP1 decoupling, consistent with previous observations^31^. We conducted the same analysis for ADNPΔN and observed increased occupancy at SINEs (Figure S2B – B2_Mm1a, B2_Mm1t), but not at LTRs (IAPEz-int) or LINEs (L1MdA_I). These findings demonstrate that association with CHD4 is not required for ADNP to bind chromatin and confirm that the HP1 subunits are required to gain access to heterochromatin.

**Figure 2.**
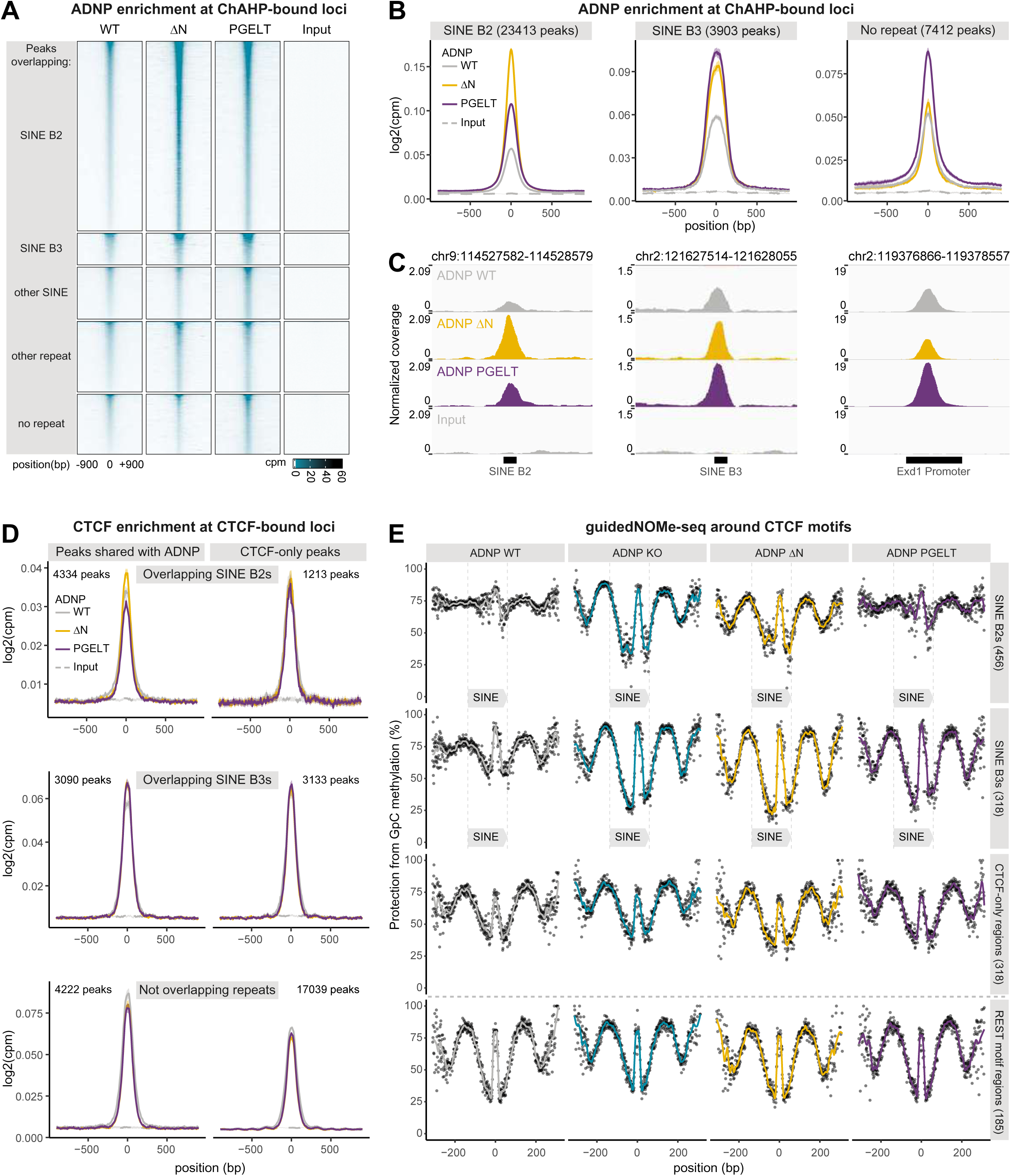
ADNPΔN binds chromatin, but fails to efficiently antagonize CTCF. **(A)** Heatmaps of ChIP-seq counts normalized to library size centered on summits of ADNP peaks (mean, n = 2 for WT, n = 3 for mutants). **(B)** Metaplots of FLAG (ADNP) ChIP-seq counts normalized to library size centered at ADNP peaks, faceted by type of genomic region they overlap (mean ± CI, n = 2 for WT, n = 3 for mutants). **(C)** IGV genome browser shots at select regions for ADNP ChIPs and input as indicated. **(D)** Metaplots of CTCF ChIP-seq counts normalized to library size centered at CTCF peaks, faceted by overlap with ADNP peaks and type of genomic region (mean ± CI, n = 2). **(E)** Protection from methylation per GpC position relative to CTCF motif in regions analyzed by guidedNOMe-seq, averaged across all fragments for each region category. The number of analyzed regions is indicated in brackets.

We have previously shown that CTCF occupancy at SINE B2 elements increases upon ablation of the ChAHP complex^30,37^. To test if this is dependent on CHD4 or HP1 subunits, we performed ChIP-seq for CTCF in the subunit-decoupled backgrounds and compared them to ADNP WT controls (Figure 2D, Figure S2C-D). Globally, CTCF occupancy was not strongly affected upon CHD4 or HP1 decoupling. However, we observed higher CTCF occupancy at SINE B2s and SINE B3s when CHD4 was decoupled. This effect was also apparent on SINE B3s, but not on SINE B2s, when HP1s were absent from the complex. Conversely, slightly less CTCF binding was evident at non-repeat regions for both mutants. Finally, CTCF occupancy at ADNP-unbound CTCF motifs was unaffected by either mutation.

To strengthen these analyses, we used the guidedNOMe-seq method to probe nucleosome positioning around CTCF motifs, which becomes more pronounced as CTCF occupancy increases (Figure 2E)^37^. CTCF and ADNP ChIP signal at the analyzed regions matched the effects seen genome-wide for all tested backgrounds, indicating that the chosen regions are representative (Figure S2E). Decoupling CHD4 from ChAHP results in prominent and symmetric protection from methylation around CTCF motifs overlapping ADNP-bound SINE B2s and B3s, with a size matching nucleosome footprints. This effect is similar to the ADNP KO condition, and specific to ADNP and CTCF co-bound regions, while CTCF sites devoid of ADNP were unaffected. Consistent with the CTCF ChIP analyses, decoupling HP1 from the complex resulted in an increase in positioning only at SINE B3s. Lastly, neither of the mutations induced any change at REST-bound regions. Together, these results show that ADNP-bound CHD4 is required for normal nucleosome distribution at ChAHP-bound loci, hinting that chromatin remodeling activity supports CTCF antagonism.

### ChAHP complex-specific inactivation of CHD4

Because CHD4 functions within multiple chromatin remodeling complexes, globally inactivating its ATPase activity obscures complex-specific roles^14,31^. To selectively abrogate ChAHP’s chromatin remodeling activity while preserving complex integrity, we generated cell lines wherein the coding sequence of CHD4 was N-terminally fused to the ADNP open reading frame at the endogenous locus. This was done in an ADNP-Avi-3xFLAG background, inserting either WT or catalytic dead CHD4 (E867Q; Walker B motif mutation), yielding a tagged in-frame chimeric transcript encoding both CHD4 and ADNP (Figure 3A). To suppress any potential association with endogenous CHD4 protein, we also fused both WT and catalytic dead CHD4 sequences to ADNPΔN. All CHD4-ADNP fusion proteins were expressed comparably to unfused ADNP-Avi-3xFLAG-, and similarly to each other (Figure S3A, Input). By co-IP, we could confirm that none of the chimeras persistently interact with unfused endogenous CHD4, which can be resolved from the chimeras by molecular weight (Figure S3A, IP). We further assessed whether the rest of the interactome changes between chimeric and unfused ADNP by proteomics (Figure S3B-C). This also allowed us to more accurately assess ADNP-CHD4 stoichiometry, which did not significantly differ between any of the chimeras and the unfused control (Figure S3B). Interactome-wide, there were no differentially interacting proteins between any of the chimeras and the unfused controls, apart from three variable background artefacts (Figure S3C). This excludes conceivable gain-of-function phenomena, such as formation of unnatural ChAHP-NuRD supercomplexes.

**Figure 3.**
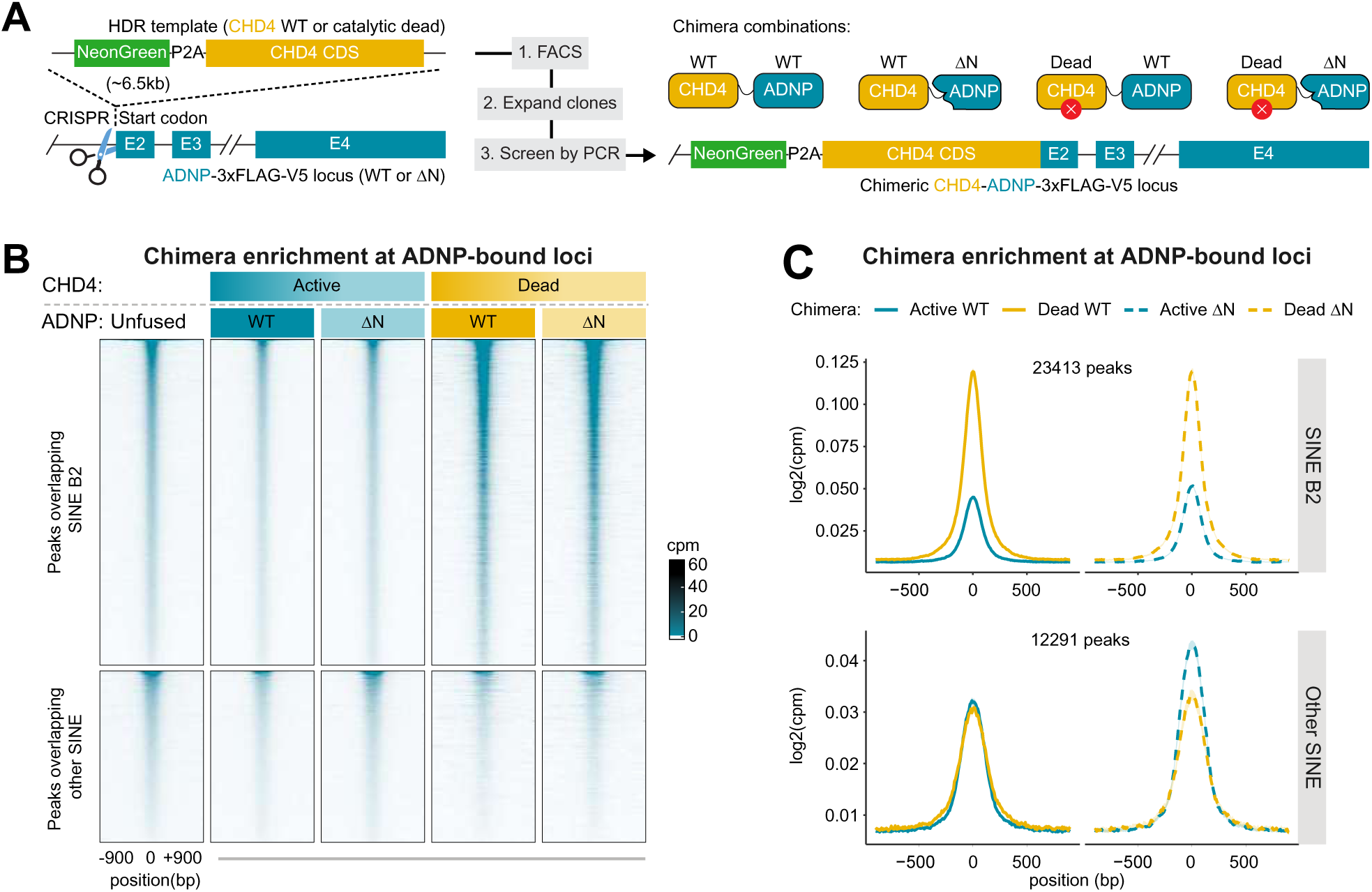
Catalytic activity of ChAHP-associated CHD4 is not required for chromatin binding. **(A)** Schematic of experimental workflow to generate active and catalytically dead (CHD4 E867Q) CHD4-ADNP chimeras. The NeonGreen fluorescent tag is separated from CHD4 with a P2A sequence, leaving 1 proline at the N-terminus of CHD4 after the ribosome skipping event. A 22 amino-acid glycine-serine linker was inserted between CHD4 and ADNP. **(B)** Heatmaps of ChIP-seq counts normalized to library size centered on summits of ADNP peaks (mean, n = 4 for Unfused ADNP, active WT and active ΔN chimeras, n = 2 for others). **(C)** Metaplots of ChIP-seq counts normalized to library size centered at ADNP peaks, faceted by type of genomic region they overlap (mean ± CI, replicates as in panel B).

### Catalytically inactive ChAHP is trapped on SINE B2s

We first used this system to assess whether CHD4’s catalytic activity within the ChAHP complex is required for chromatin engagement. To that end, we performed ChIP sequencing, leveraging the affinity tag endogenously encoded at the C-terminus of the CHD4-ADNP chimeras (Figure 3B-C). In comparison to the unfused control, ChIP enrichments at ChAHP binding sites were overall slightly reduced for chimeras harboring WT CHD4 (Figure 3B). In contrast, chromatin occupancy increased for catalytically inactive chimeric assemblies, specifically at peaks overlapping SINE B2s (Figure 3B-C). This increase was not evident at peaks overlapping the more divergent and transcriptionally inert SINE B3 elements, nor at the evolutionarily distinct SINE B1 elements (Figure 3B-C, Other SINE). These findings indicate that CHD4’s ATPase activity is dispensable for ChAHP chromatin engagement, but its absence may result in retention of the complex at specific genomic loci.

### Remodeling Activity of ChAHP counteracts CTCF binding to SINE B2s

Since the CHD4-ADNP fusion retains chromatin binding independent of CHD4’s catalytic activity, we had a tenable system to discriminate whether ChAHP competes with CTCF at SINE B2 elements through direct steric hindrance or dependent on remodeling activity. To that end, we performed CTCF ChIP-seq in cells expressing each CHD4-ADNP chimera variant (Figure 4A, Figure S4A). We observe no change in CTCF ChIP signal between CHD4^WT^-ADNP^WT^ and CHD4^WT^-ADNP^ΔN^ chimeras, indicating that the ADNP N-terminus does not influence CTCF antagonism *per se*. Conversely, cells expressing catalytically inactive ChAHP chimeras in either configuration exhibited increased CTCF occupancy at SINE B2 elements. This effect was locationally specific, as the increase was not evident beyond SINE B2s or outside ChAHP target regions. These data demonstrate that remodeling activity of ChAHP, not merely its presence, is necessary to efficiently antagonize CTCF binding at SINE B2s.

**Figure 4.**
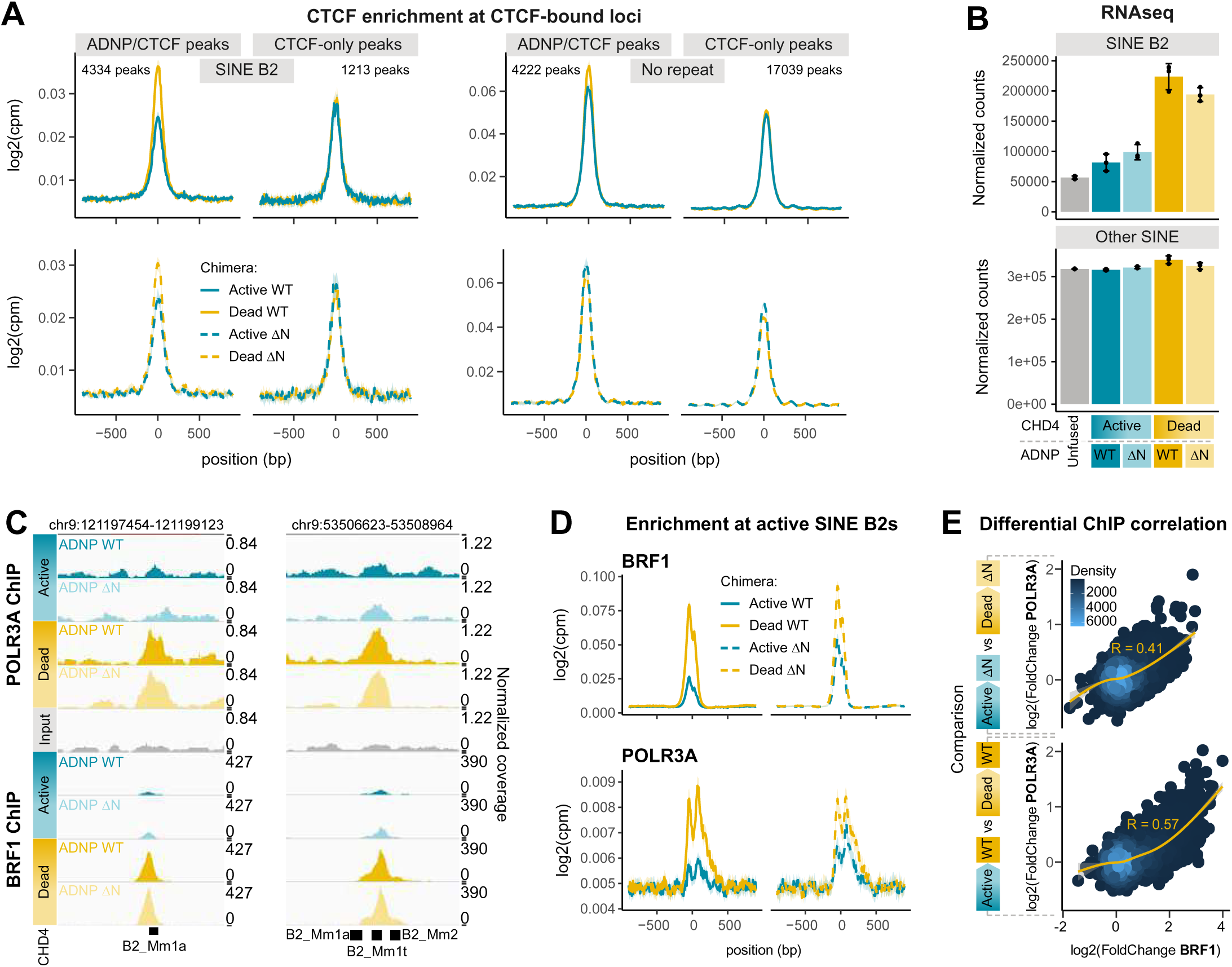
Catalytic activity of ChAHP-associated CHD4 is required to repress SINE B2 elements and antagonize CTCF. **(A)** Metaplots of CTCF ChIP-seq counts normalized to library size centered at CTCF peaks, faceted by overlap with ADNP peaks and type of genomic region (mean ± CI, n = 2). **(B)** RNA-seq reads normalized to library size for SINE B2s – combining Mm1a, Mm1t, Mm2 subfamilies – and all other SINEs (mean ± SD, n=3). **(C)** IGV genome browser shots at select regions for BRF1 and POLR3A ChIPs, and input as indicated. **(D)** Metaplots of BRF1 and POLR3A ChIP-seq counts normalized to library size centered at starts of active SINE B2 elements – Mm1a, Mm1t subfamilies (mean ± CI, n = 2). **(E)** Correlation of ChIP-seq count ratios for BRF1 and POLR3A between catalytic-dead chimeras and their corresponding catalytically active controls.

### ChAHP remodeling activity silences transcription of SINE B2s

Next, we assessed the impact of ChAHP remodeling activity on transcription of SINE B2 transposons, comparing each mutated CHD4-ADNP chimera to the CHD4^WT^-ADNP^WT^ control. Introduction of chimeras containing catalytically inactive CHD4 led to strong and significant derepression of SINE B2 elements compared to the CHD4^WT^-ADNP^WT^ control, while other SINE elements were unaffected (Figure 4B, Figure S4C). We additionally observed minor differences in LTR expression, but these were not consistent between ADNP WT- and ΔN-containing chimeras. Furthermore, major active murine LTR elements such as IAPEz and MMERVK10C, were not significantly affected (Figure S4B, Figure S4C).

These results demonstrate that transcription of SINE B2 elements by RNA polymerase III is counteracted through ChAHP remodeling activity, and not through action as a transcriptional barrier. In a related preprint, we show that ChAHP specifically prevents the transcription factor complex TFIIIB from associating with the SINE B2 promoter (Schnabl-Baumgartner *et al*. 2025. ChAHP Silences SINE Retrotransposons by Inhibiting TFIIIB Recruitment. *bioRxiv*, doi: https://doi.org/10.1101/2025.07.02.662776). To determine whether this exclusion is mediated by ChAHP’s catalytic activity, we performed ChIP-seq for the TFIIIB subunit BRF1 (Figure 4C-D). Indeed, we observed increased BRF1 binding at SINE B2 promoters in cells expressing catalytically dead ADNP-CHD4 chimeras (Figure 4C-D). Consistent with elevated transcription, the RNA polymerase III catalytic subunit POLR3A was also more enriched at the same loci. The increase in BRF1 association with SINE B2s was accompanied by a corresponding accumulation of POLR3A genome-wide, and the magnitudes of these increases strongly correlated (Figure 4E). Collectively, these data demonstrate that ChAHP represses SINE B2 transcription by preventing the recruitment of TFIIIB through a process dependent on remodeling activity.

## DISCUSSION

Since the discovery of the ChAHP complex, its mode of action has remained a subject of speculation, which included potential recruitment of auxiliary subunits and steric competition with transcription factors such as CTCF. In particular, the role of CHD4 within ChAHP had been elusive. Here, we provide direct evidence that the chromatin remodeling activity of CHD4 is indispensable for ChAHP function, but contrary to common expectation, ChAHP does not require CHD4’s ATPase activity to access its target sites. Instead, this activity is critical to counteract the chromatin binding of other transcription factors, namely CTCF and TFIIIB. This suggests that ChAHP engages with chromatin entirely independent of chromatin remodeling, or, alternatively, relies on a remodeling function that is not an integral part of the complex. Such a unidirectional dependence of a remodeler (CHD4) on a transcription factor (ADNP) is conceptually distinct from previously described TF-remodeler relationships, including labile (e.g., CTCF-SNF2H, REST-BRG1) and stable (e.g., YY1-INO80) assemblies.

Our finding that catalytically inactive ChAHP is trapped on specific chromatin loci argues against the parsimonious model in which the ChAHP complex sterically blocks transcription factor access. That is, catalytically inactive ChAHP does not support SINE B2 repression or CTCF antagonism, despite increased ChAHP occupancy specifically at these elements. This strongly suggests that direct binding competition between ChAHP and CTCF or TFIIIB makes little contribution to ChAHP function. We therefore surmise that the dominant mode of action is active nucleosome remodeling at its target sites. Altering the position or density of nucleosomes relative to key sequence motifs might occlude CTCF and TFIIIB binding sites, consistent with the general principle that nucleosomes bar TF access to DNA. This property has been demonstrated for CTCF, whose ingress to chromatin is supported by the remodeler SNF2H^18^. ChAHP may antagonize this process by disrupting nucleosome arrangements favorable for CTCF binding, thus driving a contraction in CTCF residency time on chromatin. Although it remains to be established whether TFIIIB is excluded from nucleosome-bound DNA in cells, *in vitro* work suggests RNA Pol III transcription is sensitive to nucleosome positioning^38–40^. We therefore speculate ChAHP interferes with TFIIIB function by increasing nucleosome density at TFIIIB recruitment sites.

In conclusion, we propose that ChAHP represents a functional module that enables sequence-specific targeting of chromatin remodeling activity to counteract the binding of other chromatin regulators.

### Limitations of the study

While we favor a model in which ChAHP remodels nucleosomes, an alternative explanation for transcription factor competition is that remodeling by the ChAHP complex directly ejects CTCF or TFIIIB from chromatin without repositioning a nucleosome at the TF binding site^41^. However, such a mechanism would be reliant on saturating concentrations of ChAHP and continuous occupancy at target sites to prevent CTCF/TFIIIB rebinding. Although the weak correlation between repressive potency and ChAHP occupancy at chromatin argues against this model, we cannot definitively exclude it. Since dissecting these chromatin events in more detail by cell-based steady state assays is not tenable, direct biochemical and structural evidence will be required to distinguish between these models.

## Supporting information

Cell lines used in this study

Oligonucleotide and gBlock sequences used for gRNA cloning and as homology donors for CRISPR-Cas9 editing

plasmids used in this study

Plasmid Genebank files

## ACKNOWLEDGEMENTS

We thank the members of the Bühler lab for their constant support and discussions. Special thanks to Yukiko Shimada for technical support. We thank Hubertus Kohler for cell sorting and the members of the FMI genomics and proteomics platforms for their continuous support. We are grateful to Jakob Schnabl-Baumgartner, Nazerke Atinbayeva, Fabian Pötz, and Arjun Udupa for their comments on the manuscript. This work was supported by the Novartis Research Foundation and the Swiss National Science Foundation (SNSF; grant 310030_188835).

## Author contributions

J.A. designed and performed most experiments, analyzed data, and generated the figures. F.M. generated cell lines, performed ChIP-seq and advised experiments. M.S. performed bioinformatics analyses. L.K. generated cell lines and performed guidedNOMe-seq. J.S. generated cell lines. D.H. analyzed proteomics data. E.P.F.M. prepared RNA-seq libraries and performed genomics QC. M.B. conceived and supervised the study, and secured funding. J.A., F.M. and M.B. wrote the manuscript. All authors discussed the results and commented on the manuscript.

## Declaration of generative AI and AI-assisted technologies in the writing process

During the preparation of this work the authors used ChatGPT 4o to improve the readability and language of the manuscript. After using this tool, the authors reviewed and edited the content as needed and take full responsibility for the content of the published article.

## STAR METHODS

### RESOURCE AVAILABILITY

#### Lead contact

Further information and requests for resources and reagents should be directed to and will be fulfilled by the lead contact, Marc Bühler.

### Materials availability

Cell lines and plasmids are available upon request.

### Data and code availability

Sequencing data from ChIP-seq and mRNA-seq have been deposited at GEO (accession number GSE297864) and are publicly available as of the date of publication. The mass spectrometry proteomics data have been deposited to the ProteomeXchange Consortium via the PRIDE^42^ partner repository with the dataset identifier PXD064364 and PXD064333. This paper analyzes existing, publicly available data. Their accession numbers are listed in the key resources table.

All original code has been deposited on GitHub and is publicly available at https://github.com/xxxmichixxx/ADNP_CHD4

## EXPERIMENTAL MODEL AND SUBJECT DETAILS

Mouse embryonic stem cells (129 × C57BL/6 background) with BirA and Cre insertions in the Rosa26 locus^43,44^ were cultured on gelatin-coated dishes in ES medium containing DMEM (Sigma D5648), supplemented with 15% fetal bovine serum (FBS; GIBCO), 1 × non-essential amino acids (GIBCO), 1 mM sodium pyruvate (GIBCO), 2 mM l-glutamine (GIBCO), 0.1 mM 2-mercaptoethanol (Sigma), 50 mg/ml penicillin, 80 mg/ml streptomycin, 3 μM glycogen synthase kinase 3β (GSK) inhibitor (Sigma, CHIR99021), 10 μM MEK inhibitor (Tocris, PD0325901), and homemade LIF, at 37°C in 5% CO2.

## METHOD DETAILS

### Genome editing

Cells were trypsinized, counted, seeded, and immediately transfected using lipofectamine 3000 (Invitrogen) according to manufacturer instructions. Generally, 300k cells were seeded in 6-well plates and transfected with a total of 1-1.5mg DNA. For endogenous tagging, a plasmid harboring a desired homology repair cassette was included. The cells were then trypsinized, counted and 15000 cells were seeded on a 10cm dish for colony formation without puromycin selection. When colonies were sufficiently large (4-8 days), they were manually picked and split into two 96-well plates for screening and expansion. ADNP KO cells were made as previously described^27,30,31^. For CHD4-ADNP fusion cell lines, single cells were sorted by FACS into 96-well plates 3-5 days after transfection, selected by matching fluorescence intensity to a homozygously-tagged ADNP-NeonGreen-3xFLAG mESC cell line. 10μM ROCK inhibitor (Y-27632) was included in the media during the first 24h after sorting. Screening for both positive editing events and unedited loci was performed by PCR and Sanger sequencing of the products.

### Western blotting

Protein samples in lauryl-dodecyl sulfate sample buffer (LDS) were separated by standard SDS-PAGE on 4-12% Bis-Tris gradient gels (Novex Bolt, Invitrogen) or 3-8% Tris-acetate gradient gels (NuPage, Invitrogen). Separated proteins were transferred onto PVDF membranes (Milipore) in transfer buffer (Bjerrum-Schaeffer-Nielsen + 0.4% SDS) using a semi-dry transfer procedure (TransBlot Turbo, BioRad) with transfer parameters: 1.3A, 25V, 12min for one gel. The membranes were blocked in 3-5% skimmed milk (Sigma) dissolved in Tris buffered saline + 0.2% v/v Tween-20 (TBST). Antibody incubations were performed with antibodies diluted in blocking solution to empirically determined concentrations for a minimum of 45min up to a maximum of 24h. When re-probing, membranes were first treated with 0.01% NaN_3_ for 30-60min to quench the HRP from previous staining rounds. Between each antibody incubation, the membranes were washed for a minimum of 45min in TBST with at least 4 buffer exchanges. The horseradish peroxidase system (Immobilon, Milipore), coupled to camera-based detection (AI600 or AI680, Agilent technologies) was used to visualize protein bands.

### Immune precipitations

Cells from one 10cm dish per condition were harvested by trypsinization before centrifugation (200g_av_, 5min). Cells were washed once in room temperature PBS and snap frozen or immediately taken forward for lysis. Pellets were lysed in NP-40 lysis buffer supplemented with protease inhibitors and benzonase (20mM Tris-HCl pH7.4, 150mM NaCl, 1% (v/v) NP-40, 0.1% (v/v) sodium deoxycholate,1xHALT protease inhibitor cocktail, 200U Turbo Benzonase (Milipore)) for 20min at 12°C and cleared by centrifugation at 4°C (16000g_av_, 20min). Protein concentration in the lysates was determined by Bradford assay against BSA and 1-3mg protein-lysate equivalent was loaded onto 10μL of bead slurry (Dynabeads, GE healthcare) prewashed twice with lysis buffer. For FLAG IPs, 2μg of antibody was added per 1mg of protein. For Strep IP Streptavidin M280 dynabeads (Invitrogen) were used. Beads were incubated with lysate for 2h at 4°C, washed twice with lysis buffer and twice with wash buffer (20mM Tris-HCl pH7.4, 150mM NaCl, 0.1%(v/v) NP-40) before addition of LDS. All bead separation steps were done using magnetic racks.

### Proteomics

For IP-MS, immune precipitations were performed as normal with the addition of 2 washing steps without detergent (20mM Tris pH7.5, 150mM NaCl), followed by on-bead digestion. Beads were resuspended in 5 µL of digestion buffer (3M GuaHCl, 20mM EPPS pH8.5, 10mM CAA, 5mM TCEP) and 1 µL of 0.2 mg/mL LysC protease (Promega) in 50 mM HEPES (pH 8.5) was added. Proteins were digested for 2 h rotating at room temperature. The samples were diluted with 17 µL of 50 mM HEPES (pH 8.5) and digested with 1 µL of 0.2 mg/mL trypsin (Promega) in 0.2 mM HCl at 37°C with interval mixing at 2000 RPM for 30 s every 15 min.

Digested peptides were acidified with 0.8% TFA (final) and analyzed by LC–MS/MS as follows:

For **Figure S1E**:

Instrument: Orbitrap LUMOS (Thermo Fisher Scientific) with VanquishNeo-nLC and an easy source with a 75umx15cm EasyC18 column was used. The samples were loaded on a C18 (0.3x5mm) trap and backward flush was used for the analysis.

Gradient 80min: 0-1min 2-6%B in A, 1-43min 6-20%, 43-65.5min 20-35%, 65.5-67.5min 35-45%, 67.5-68.5min 45-100%, 68.5-80min 100%. Buffer A: 0.1%FA in H2O; Buffer B: 0.1%FA, 80% MeCN in H2O, at RT and the flow rate during the gradient was 200ul/min.

For **Figure S3B-C**:

Instrument: Orbitrap LUMOS (Thermo Fisher Scientific) with VanquishNeo-nLC and an easy source with a 75umx15cm EasyC18 column was used. The samples were loaded on a C18 (0.3x5mm) trap and backward flush was used for the analysis.

Gradient 80min: 0-1min 2-6%B in A, 1-43min 6-20%, 43-65.5min 20-35%, 65.5-67.5min 35-45%, 67.5-68.5min 45-100%, 68.5-80min 100%. Buffer A: 0.1%FA in H2O; Buffer B: 0.1%FA, 80% MeCN in H2O, at RT and the flow rate during the gradient was 200ul/min.

Data were acquired using 120,000 resolution for the peptide measurements in the Orbitrap and a top T (3-sec) method with HCD fragmentation for each precursor and fragment measurement in the ion trap following the manufacturer guidelines (Thermo Scientific).

### ChIP-seq

ChIP experiments were performed with at least 2 different clones from endogenously tagged cell lines. For ADNP, CHD4-ADNP chimeras, and BRF1, the ChIPs were performed using antibodies against the affinity tag (3xFLAG), and for CTCF and POLR3A using specific antibodies as detailed further below. Harvesting was performed by trypsinization, and cells were counted for each sample. For each ChIP 10^7^ cells were collected. The cells were crosslinked for 8 min at room temperature in 10ml PBS supplemented with 1% formaldehyde (Sigma, F8775). Cross-linking was quenched by adding glycine to a final concentration of 0.125 mM and incubating at room temperature for 1min, and on ice for 3min. Cells were pelleted by centrifugation at 500 g for 3 min at 4°C and the pellet was lysed in 10mL lysis buffer A (50 mM HEPES pH 8.0, 140 mM NaCl, 1 mM EDTA, 10% glycerol, 0.5% NP40, 0.25% Triton X-100) for 10 minutes on ice. After centrifugation the pellet was resuspended in 10 mL buffer B (10 mM Tris pH 8, 1 mM EDTA, 0.5 mM EGTA and 200 mM NaCl) and incubated for on ice for 5min. The samples were centrifuged at 500 g for 3 min at 4°C, and the pellets lysed in 180ul buffer C (50 mM Tris pH 8, 5 mM EDTA, 1% SDS, 100 mM NaCl) for 2min at room temperature and on ice for 10 min. The lysates were diluted in 1.6 mL ice cold TE buffer and sonicated in 15 mL tubes two times 10 cycles, 30 s ON / 30 s OFF, at 4°C (Bioruptor Pico, BioRad). Then, 200ul 10x ChIP buffer (0.1% SDS, 10% Triton X-100, 12mM EDTA, 167mM Tris-HCl pH 8, 1.67M NaCl) was added, and the chromatin was transferred into 2 mL Eppendorf tubes before centrifugation for 10 min at 16000 g, 4°C. 5% sheared chromatin was reserved for the input control, while the rest was transferred into fresh tubes. Generally, beads were prebound with antibody in PBS+0.2% Tween, and washed twice with 1x ChIP buffer, before being added to sonicated chromatin. For FLAG ChIPs, 20uL Protein G Dynabeads and 3uL α-FLAG. For CTCF and POLR3A, 15uL Protein G were mixed with 15uL Protein A Dynabeads and 3uL antibody added. Samples were incubated for 4h at 4°C. ChIPs were washed for 1 minute each for each step, 4 times RIPA (10mM Tris-HCl pH 8.0, 1mM EDTA pH 8.0, 140mM NaCl, 1% Triton X-100, 0.1% SDS, 0.1% Na-deoxycholate), 2 times RIPA500 (10mM Tris-HCl pH 8.0, 1mM EDTA pH 8.0, 500mM NaCl, 1% Triton X-100, 0.1% SDS, 0.1% Na-deoxycholate), 2 times Li-wash buffer (10mM Tris-HCl pH 8.0, 1mM EDTA, pH 8.0, 250mM LiCl, 0.5% NP-40, 0.5% Na-deoxycholate), 1x TE (10mM Tris-HCl pH 8.0, 1mM EDTA). Beads were transferred to a fresh tube during the last wash and wash buffer was completely removed before adding 50uL elution buffer (10mM Tris-HCl pH 8.0, 1mM EDTA pH 8.0, 150mM NaCl, 1% SDS) and incubating 20 minutes at 65°C with constant shaking. Elution was repeated once more with 50uL elution buffer for 20 minutes and eluates were pooled, 2 ul RNaseA (20μg/ul) were added and incubated for 1 h at 37°C. Then 2ul Proteinase K (20mg/ml) was added and samples were incubated 2 h at 55°C followed by decrosslinking for 6 h at 65°C. Input samples were adjusted to 100ul total volume with elution buffer and processed equivalently to ChIP samples. DNA was purified by adding 20ul AMPure XP beads, 6 ul 5M NaCl, and 130ul Isopropanol and incubating for 10 min at RT after thorough mixing. The beads were collected on a magnetic rack, washed twice with 80% EtOH and DNA was eluted in 30ul 10mM Tris pH8.0 for 5 min at 37°C. 25ul ChIP DNA or 10ng Input DNA were used to generate libraries using the NEBNext Ultra II Library Prep Kit for Illumina (NEB). Reactions were scaled down to half otherwise processing was according to the manufacturer’s manual. Libraries were sequenced 51bp paired-end on a NovaSeq6000 instrument (Illumina).

### RNA-seq

RNA was isolated from cells using the RNeasy miniprep kit (Qiagen) according to the manufacturer’s instructions, including genomic DNA pre-filtering and DNaseI treatment using 2U of RQ1 DNaseI (Promega) for 30min. The concentration was determined with the RNA Broad Range reagents on a Qubit 2.0 system according to the manufacturer’s instruction. Libraries were prepared using the Illumina Stranded Total RNA Library Prep, including a ribosomal RNA depletion step, and sequenced on the Illumina NovaSeq 6000 (51-nt paired-end reads).

### Computational methods Structure prediction

ChAHP structure prediction was performed on the AlphaFold Server (version from October 2024)^36^, by importing the full length mouse protein sequences of CHD4, ADNP, and two HP1γ units, alongside 1 ATP molecule, 1 Mg^2+^ ion, and 13 Zn^2+^ ions. The output folder was imported into AlphaBridge^45^ for interpretation, and individual models further examined in ChimeraX^46^.

### ChIP seq alignment

Reads were mapped to the mouse mm39 genome using STAR version 2.7.3^47^ and repeat sensitive settings as suggested by Teissandier et al^48^. The parameters used were STAR: --outSAMtype BAM SortedByCoordinate --winAnchorMultimapNmax 5000 --alignEndsType EndToEnd --alignIntronMax 1 --alignMatesGapMax 2000 -- seedSearchStartLmax 30 --alignTranscriptsPerReadNmax 100000 -- alignWindowsPerReadNmax 100000 --alignTranscriptsPerWindowNmax 300 -- seedPerReadNmax 1000 --seedPerWindowNmax 300 --seedNoneLociPerWindow 1000 --outBAMsortingBinsN 100 --clip3pAdapterSeq CTGTCTCTTATACACATCT AGATGTGTATAAGAGACAG --limitBAMsortRAM 10000000000 -- alignSJDBoverhangMin 999.

### Peak finding

Peak calling was performed using MACS3 version 3.0.1^49^ on combined replicates for each ChIP condition. Read counts within peaks (resized to 300 bp around peak summits) were calculated for both ChIP and Input samples using featureCounts() from the Rsubread package^50^. Counts were normalized to library size (counts per million, CPM), and peaks were retained if they showed >1.2-fold ChIP/Input enrichment and CPM >1 in all replicates.

### ChIP-seq coverage plots

Representative ChIP-seq tracks were generated by merging replicate bam files, and converting them to the bigwig format. Tracks were visualized using the IGV genome browser software (version 2.8.6)^51^, and arranged for display in Adobe Illustrator (29.0.1).

### ChIP-seq heatmap generation

ChIP-seq heatmaps were generated using the MiniChip R package (version 0.0.0.9600). The SummitHeatmap() function was used with parameters: span = 900, step = 9, minOverlap = 1, useCPM = TRUE, PairedEnd = TRUE, minMQS = 0, strand = 0, readExtension3 = 0, readShiftSize = "halfInsert", read2pos = 5, mode = "Q". Heatmaps were drawn using DrawSummitHeatmaps() with parameters: use.log = FALSE, medianCpm = medianCpm, topCpm = topCpm, TargetHeight = 1000, orderSample = 1, orderWindows = 15, summarizing = "mean", show_axis = FALSE.

### ChIP-seq metaplots

To generate metaplots we used the CumulativePlots() function of the MiniChip R package(version 0.0.0.9600).

### RNA-seq read mapping and quantification

RNA-seq reads were aligned to the mouse genome (GRCm39) using STAR aligner with following parameters: --outFilterType BySJout --outFilterMultimapNmax 100000 -- outFilterMismatchNmax 3 --winAnchorMultimapNmax 20000 --alignMatesGapMax 350 - -seedSearchStartLmax 30 alignTranscriptsPerReadNmax 30000 -- alignWindowsPerReadNmax 30000 --alignTranscriptsPerWindowNmax 300 -- seedPerReadNmax 3000 --seedPerWindowNmax 300 --seedNoneLociPerWindow 1000--outSAMattributes NH HI NM MD AS nM --outMultimapperOrder Random -- outSAMmultNmax 1 --outSAMunmapped Within --limitBAMsortRAM 10000000000.

Uniquely mapped reads were quantified at the gene level using featureCounts with the GENCODE vM34 annotation.

For repetitive element analysis, a custom annotation file was created from RepeatMasker data (GRCm39.primary_assembly.repMasker_sensitive.bed). including multimapping reads. Read counts were generated using featureCounts with settings allowing for multiple mapping reads, multiple overlaps, and no read fraction assignment. Simple repeats, low complexity regions, and rRNA were excluded from the analysis to focus on biologically relevant elements. Only repeats with at least 100 normalized read counts summed across samples were included in downstream analyses.

### Differential expression analysis

Differential expression analysis was performed using DESeq2^52^ for both genes and repetitive elements. For differential expression significance, a fold change threshold of 2-fold (|log₂FC| > 1) and FDR < 0.05 were used. Expression changes were evaluated at each condition compared to the respective controls as indicated in the figures.

For quantitative analysis of specific repeat families, normalized count data was extracted from the DESeq2 object to allow direct comparison between conditions. Count normalization was performed using DESeq2’s variance stabilizing transformation. Mean normalized counts with standard deviation were calculated for each experimental condition across biological replicates.

### MA-plots

RNA-seq data visualization was performed using R (version 4.1.0) with the tidyverse, and ggplot2 packages. Differential expression results were visualized using MA plots, with significantly regulated genes and repeat elements highlighted (|log₂FC| > 1 and FDR < 0.05). For repeat elements, specific families were color-coded by class.

### Proteomics

Peptides were identified with MaxQuant version 2.2.0.0 using the search engine Andromeda^53^ or using FragPipe version 22.0^54^. The mouse subset of the UniProt version 2023_03 or 2023_08 combined with the contaminant DB from MaxQuant was searched and the protein and peptide FDR values were set to 0.05. Statistical analysis was done using limma within the einProt package (version 0.9.6)^55^. Results were filtered to remove reverse hits, contaminants and peptides found in only one sample. Missing values were imputed and potential interactors visualized in volcano plots.

## SUPPLEMENTAL INFORMATION

Document S1. Figures S1-S4

Table S1. mESC lines used in this study, related to STAR Methods

Table S2. Oligonucleotides used in this study, related to STAR Methods

Table S3. Plasmids used in this study, related to STAR Methods

Data S1. Annotated GenBank files for all plasmids used in this study, related to STAR Methods

**Figure S1.**
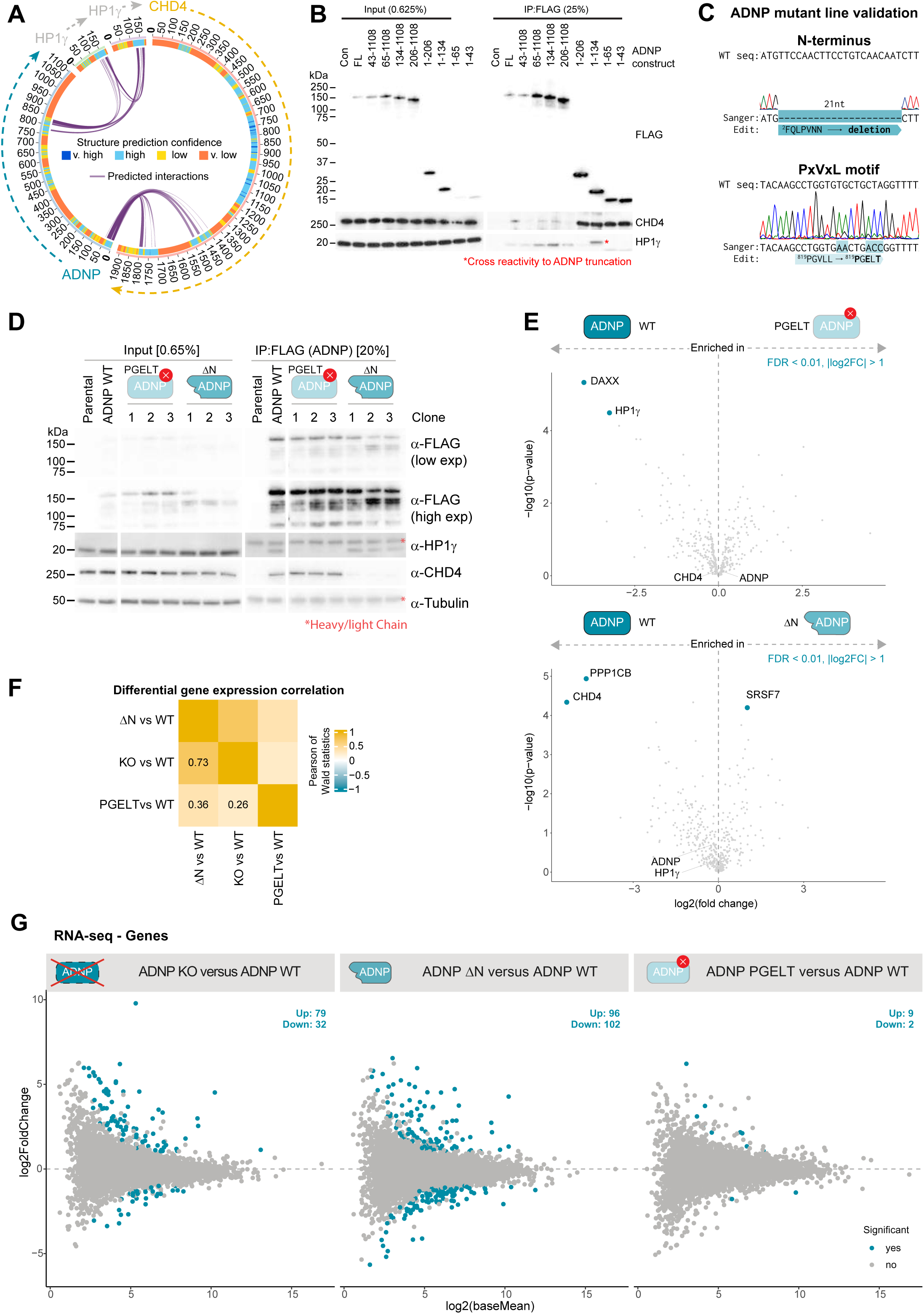
Establishment and characterization of ChAHP subunit-decoupled cell lines, related to Figure 1. **(A)** Predicted interaction surfaces between ChAHP subunits based on AlphaFold3 modeling, inferred using the AlphaBridge tool^45^. **(B)** mESCs were transiently transfected with plasmids encoding 3xFLAG-Avi tagged fragments of ADNP as indicated, subjected to co-IP with α-FLAG coated beads, and analyzed by western blotting. **(C)** Example sequence traces of endogenously edited cell lines with mutations in either the CHD4-interacting interface (N-terminus) or the HP1-interacting site (PxVxL motif) of ADNP. **(D-E)** Endogenously edited subunit-decoupled cell lines were subjected to co-IP against FLAG and analyzed by western blotting (D) and proteomics (E). For (E), colored dots represent significantly changing interactors between each mutant and the WT control. PPP1CB and SRSF7 were detected in the negative control samples, and therefore represent background variation between samples, rather than a biologically relevant difference (see dataset on PRIDE). **(F-G)** Differential genes expression analysis comparing WT cells to ADNP KO cells or cells encoding subunit-decoupled ADNP (n=2 for PGELT mutant data, n=3 for others), including a correlation of gene expression changes (F) and MA plots (G). In (G), significant hits are highlighted in blue, and the numbers given in each MA plot (FDR < 0.05, |log2FoldChange| > 1).

**Figure S2.**
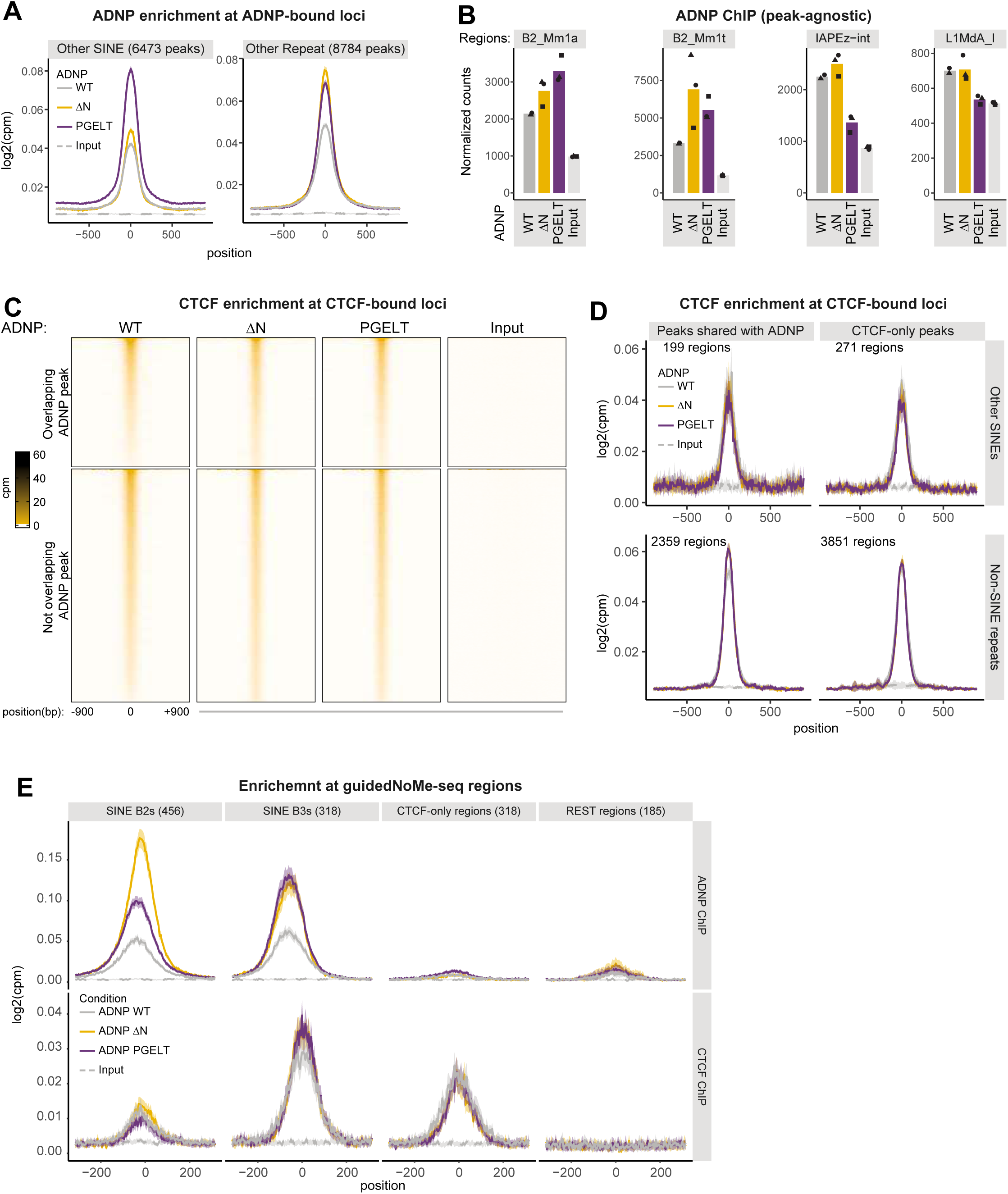
ADNPΔN binds chromatin, but fails to efficiently antagonize CTCF, related to Figure 2. **(A)** Metaplots of FLAG (ADNP) ChIP-seq counts normalized to library size centered at ADNP peaks, faceted by type of genomic region they overlap (mean ± CI, n = 2 for WT, n = 3 for mutants) – additional region types supporting Figure 2B. **(B)** Summed reads over repeat annotations normalized to library size for input and a series of ChIP samples as indicated (mean with individual replicates shown, n=2 for ADNP WT ChIP, n=3 for others). **(C)** Heatmaps of ChIP-seq counts normalized to library size centered on summits of CTCF peaks (mean, n = 2). **(D)** Metaplots of CTCF ChIP-seq counts normalized to library size centered at CTCF peaks, faceted by overlap with ADNP peaks and type of genomic region (mean ± CI, n = 2). **(E)** Same as (D), but specifically for regions analyzed by guidedNOMe-seq.

**Figure S3.**
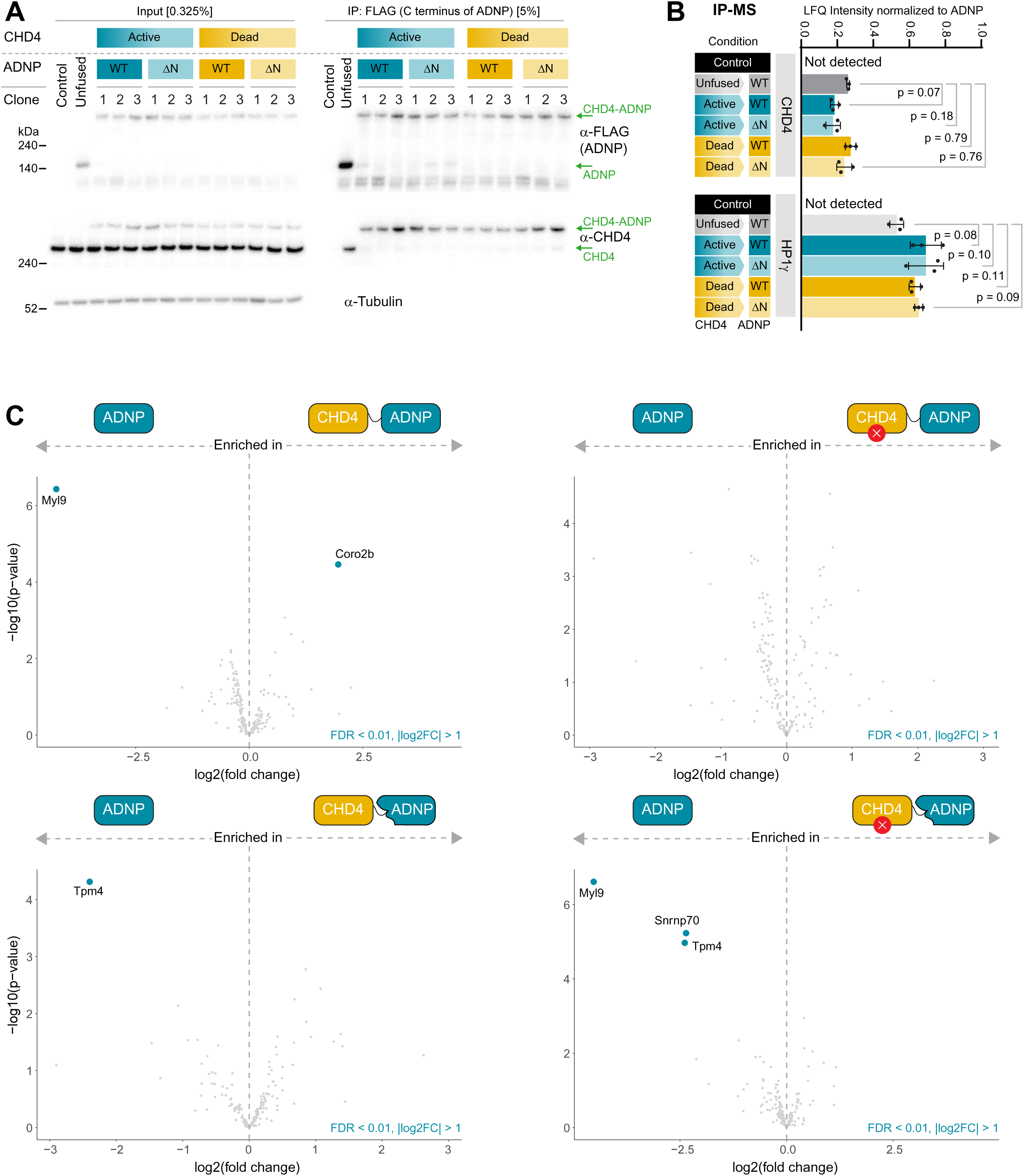
Catalytic activity of ChAHP-associated CHD4 is not required for chromatin binding, related to Figure 3. **(A-C)** Cell lines edited to express CHD4-ADNP chimeras were subjected to immune precipitation against FLAG, followed by western blotting (A) and proteomics (B-C) analyses. In (A), the corresponding sizes for natural CHD4 or ADNP proteins versus the chimeric assemblies are highlighted. For (B), label-free quantification of CHD4 and HP1γ proteins in ADNP IPs are shown, normalized to amount of ADNP (mean ± SD, n = 3). In (C), colored dots represent significantly changing interactors between each variant and the double WT control (FDR < 0.01, |log2FC| > 1).

**Figure S4.**
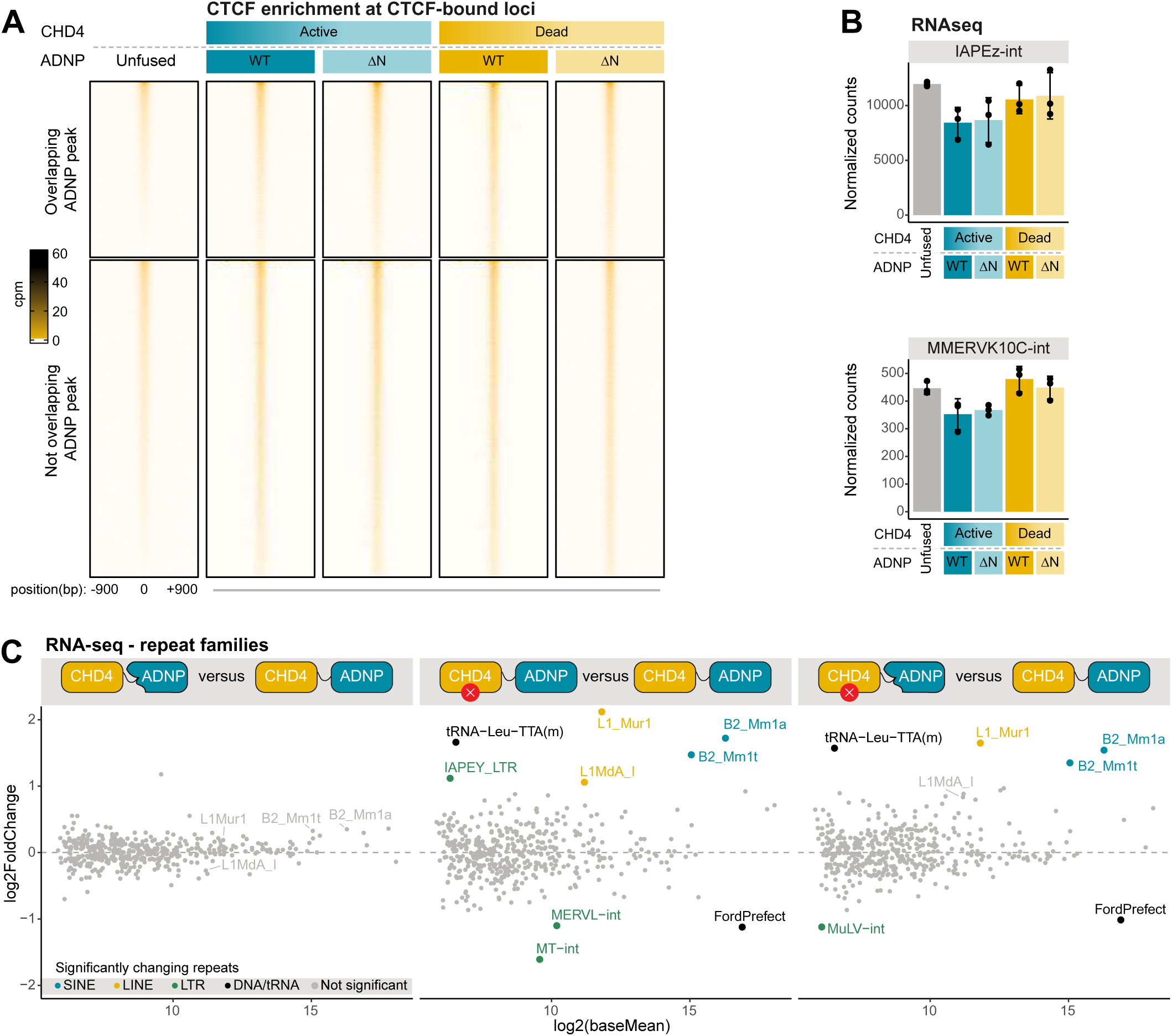
Catalytic activity of ChAHP-associated CHD4 is required to repress SINE B2 elements and antagonize CTCF, related to Figure 4. **(A)** Heatmaps of ChIP-seq counts normalized to library size centered on summits of CTCF peaks (mean, n = 2). **(B)** RNA-seq reads normalized to library size for representative LTR retrotransposons (mean ± SD, n=3). **(C)** Differential expression analysis for repeat families between CHD4-ADNP chimera variants and the double WT chimera control (n=3). Significant hits are highlighted with colors (FDR < 0.05, |log2FoldChange| > 1). No repeat family is significantly differentially expressed between chimeras encoding active CHD4.

## REFERENCES

1. Wolf, D., and Goff, S.P. (2009). Embryonic stem cells use ZFP809 to silence retroviral DNAs. Nature 458, 1201–1204. 10.1038/nature07844.

2. Imbeault, M., Helleboid, P.-Y., and Trono, D. (2017). KRAB zinc-finger proteins contribute to the evolution of gene regulatory networks. Nature 543, 550–554. 10.1038/nature21683.

3. Ecco, G., Imbeault, M., and Trono, D. (2017). KRAB zinc finger proteins. Development 144, 2719–2729. 10.1242/dev.132605.

4. Schultz, D.C., Friedman, J.R., and Rauscher, F.J. (2001). Targeting histone deacetylase complexes via KRAB-zinc finger proteins: the PHD and bromodomains of KAP-1 form a cooperative unit that recruits a novel isoform of the Mi-2α subunit of NuRD. Genes Dev. 15, 428–443. 10.1101/gad.869501.

5. Smallwood, A., Estève, P.-O., Pradhan, S., and Carey, M. (2007). Functional cooperation between HP1 and DNMT1 mediates gene silencing. Genes Dev. 21, 1169–1178. 10.1101/gad.1536807.

6. Sadic, D., Schmidt, K., Groh, S., Kondofersky, I., Ellwart, J., Fuchs, C., Theis, F.J., and Schotta, G. (2015). Atrx promotes heterochromatin formation at retrotransposons. EMBO Rep. 16, 836–850. 10.15252/embr.201439937.

7. Groh, S., Milton, A.V., Marinelli, L.K., Sickinger, C.V., Russo, A., Bollig, H., Almeida, G.P. de, Schmidt, A., Forné, I., Imhof, A., et al. (2021). Morc3 silences endogenous retroviruses by enabling Daxx-mediated histone H3.3 incorporation. Nat. Commun. 12, 5996. 10.1038/s41467-021-26288-7.

8. Sachs, P., Ding, D., Bergmaier, P., Lamp, B., Schlagheck, C., Finkernagel, F., Nist, A., Stiewe, T., and Mermoud, J.E. (2019). SMARCAD1 ATPase activity is required to silence endogenous retroviruses in embryonic stem cells. Nat. Commun. 10, 1335. 10.1038/s41467-019-09078-0.

9. Zhu, F., Farnung, L., Kaasinen, E., Sahu, B., Yin, Y., Wei, B., Dodonova, S.O., Nitta, K.R., Morgunova, E., Taipale, M., et al. (2018). The interaction landscape between transcription factors and the nucleosome. Nature 562, 76–81. 10.1038/s41586-018-0549-5.

10. Grand, R.S., Pregnolato, M., Baumgartner, L., Hoerner, L., Burger, L., and Schübeler, D. (2024). Genome access is transcription factor-specific and defined by nucleosome position. Mol. Cell 84, 3455–3468.e6. 10.1016/j.molcel.2024.08.009.

11. Kornberg, R.D. (1974). Chromatin Structure: A Repeating Unit of Histones and DNA. Science 184, 868–871. 10.1126/science.184.4139.868.

12. Michael, A.K., and Thomä, N.H. (2021). Reading the chromatinized genome. Cell 184, 3599–3611. 10.1016/j.cell.2021.05.029.

13. Lai, A.Y., and Wade, P.A. (2011). Cancer biology and NuRD: a multifaceted chromatin remodelling complex. Nat. Rev. Cancer 11, 588–596. 10.1038/nrc3091.

14. Bornelöv, S., Reynolds, N., Xenophontos, M., Gharbi, S., Johnstone, E., Floyd, R., Ralser, M., Signolet, J., Loos, R., Dietmann, S., et al. (2018). The Nucleosome Remodeling and Deacetylation Complex Modulates Chromatin Structure at Sites of Active Transcription to Fine-Tune Gene Expression. Mol. Cell 71, 56–72.e4. 10.1016/j.molcel.2018.06.003.

15. Tagami, H., Ray-Gallet, D., Almouzni, G., and Nakatani, Y. (2004). Histone H3.1 and H3.3 Complexes Mediate Nucleosome Assembly Pathways Dependent or Independent of DNA Synthesis. Cell 116, 51–61. 10.1016/s0092-8674(03)01064-x.

16. Dyer, M.A., Qadeer, Z.A., Valle-Garcia, D., and Bernstein, E. (2017). ATRX and DAXX: Mechanisms and Mutations. Cold Spring Harb. Perspect. Med. 7, a026567. 10.1101/cshperspect.a026567.

17. Xue, Y., Gibbons, R., Yan, Z., Yang, D., McDowell, T.L., Sechi, S., Qin, J., Zhou, S., Higgs, D., and Wang, W. (2003). The ATRX syndrome protein forms a chromatin-remodeling complex with Daxx and localizes in promyelocytic leukemia nuclear bodies. Proc. Natl. Acad. Sci. 100, 10635–10640. 10.1073/pnas.1937626100.

18. Barisic, D., Stadler, M.B., Iurlaro, M., and Schübeler, D. (2019). Mammalian ISWI and SWI/SNF selectively mediate binding of distinct transcription factors. Nature 569, 136–140. 10.1038/s41586-019-1115-5.

19. Wiechens, N., Singh, V., Gkikopoulos, T., Schofield, P., Rocha, S., and Owen-Hughes, T. (2016). The Chromatin Remodelling Enzymes SNF2H and SNF2L Position Nucleosomes adjacent to CTCF and Other Transcription Factors. PLoS Genet. 12, e1005940. 10.1371/journal.pgen.1005940.

20. Fryer, C.J., and Archer, T.K. (1998). Chromatin remodelling by the glucocorticoid receptor requires the BRG1 complex. Nature 393, 88–91. 10.1038/30032.

21. Krishnakumar, R., Chen, A.F., Pantovich, M.G., Danial, M., Parchem, R.J., Labosky, P.A., and Blelloch, R. (2016). FOXD3 Regulates Pluripotent Stem Cell Potential by Simultaneously Initiating and Repressing Enhancer Activity. Cell Stem Cell 18, 104–117. 10.1016/j.stem.2015.10.003.

22. Swinstead, E.E., Paakinaho, V., Presman, D.M., and Hager, G.L. (2016). Pioneer factors and ATP-dependent chromatin remodeling factors interact dynamically: A new perspective. BioEssays 38, 1150–1157. 10.1002/bies.201600137.

23. Singhal, N., Graumann, J., Wu, G., Araúzo-Bravo, M.J., Han, D.W., Greber, B., Gentile, L., Mann, M., and Schöler, H.R. (2010). Chromatin-Remodeling Components of the BAF Complex Facilitate Reprogramming. Cell 141, 943–955. 10.1016/j.cell.2010.04.037.

24. Cai, Y., Jin, J., Yao, T., Gottschalk, A.J., Swanson, S.K., Wu, S., Shi, Y., Washburn, M.P., Florens, L., Conaway, R.C., et al. (2007). YY1 functions with INO80 to activate transcription. Nat. Struct. Mol. Biol. 14, 872–874. 10.1038/nsmb1276.

25. Wu, S., Shi, Y., Mulligan, P., Gay, F., Landry, J., Liu, H., Lu, J., Qi, H.H., Wang, W., Nickoloff, J.A., et al. (2007). A YY1–INO80 complex regulates genomic stability through homologous recombination–based repair. Nat. Struct. Mol. Biol. 14, 1165–1172. 10.1038/nsmb1332.

26. Paredes, R., Gutiérrez, J., Gutierrez, S., Allison, L., Puchi, M., Imschenetzky, M., Wijnen, A. van, Lian, J., Stein, G., Stein, J., et al. (2002). Interaction of the 1α,25-dihydroxyvitamin D3 receptor at the distal promoter region of the bone-specific osteocalcin gene requires nucleosomal remodelling. Biochem. J. 363, 667–676. 10.1042/bj3630667.

27. Ostapcuk, V., Mohn, F., Carl, S.H., Basters, A., Hess, D., Iesmantavicius, V., Lampersberger, L., Flemr, M., Pandey, A., Thomä, N.H., et al. (2018). Activity-dependent neuroprotective protein recruits HP1 and CHD4 to control lineage-specifying genes. Nature 557, 739–743. 10.1038/s41586-018-0153-8.

28. Mandel, S., Rechavi, G., and Gozes, I. (2007). Activity-dependent neuroprotective protein (ADNP) differentially interacts with chromatin to regulate genes essential for embryogenesis. Dev. Biol. 303, 814–824. 10.1016/j.ydbio.2006.11.039.

29. Helsmoortel, C., Silfhout, A.T.V., Coe, B.P., Vandeweyer, G., Rooms, L., Ende, J. van den, Schuurs-Hoeijmakers, J.H.M., Marcelis, C.L., Willemsen, M.H., Vissers, L.E.L.M., et al. (2014). A SWI/SNF-related autism syndrome caused by de novo mutations in ADNP. Nat Genet 46, 380–384. 10.1038/ng.2899.

30. Kaaij, L.J.T., Mohn, F., Weide, R.H. van der, Wit, E. de, and Bühler, M. (2019). The ChAHP Complex Counteracts Chromatin Looping at CTCF Sites that Emerged from SINE Expansions in Mouse. Cell 178, 1437–1451.e14. 10.1016/j.cell.2019.08.007.

31. Ahel, J., Pandey, A., Schwaiger, M., Mohn, F., Basters, A., Kempf, G., Andriollo, A., Kaaij, L., Hess, D., and Bühler, M. (2024). ChAHP2 and ChAHP control diverse retrotransposons by complementary activities. Genes Dev. 38, 554–568. 10.1101/gad.351769.124.

32. Thorn, G.J., Clarkson, C.T., Rademacher, A., Mamayusupova, H., Schotta, G., Rippe, K., and Teif, V.B. (2022). DNA sequence-dependent formation of heterochromatin nanodomains. Nat Commun 13, 1861. 10.1038/s41467-022-29360-y.

33. Mandel, S., and Gozes, I. (2007). Activity-dependent Neuroprotective Protein Constitutes a Novel Element in the SWI/SNF Chromatin Remodeling Complex. J. Biol. Chem. 282, 34448– 34456. 10.1074/jbc.m704756200.

34. Mosch, K., Franz, H., Soeroes, S., Singh, P.B., and Fischle, W. (2011). HP1 recruits activity-dependent neuroprotective protein to H3K9me3 marked pericentromeric heterochromatin for silencing of major satellite repeats. PloS one 6, e15894. 10.1371/journal.pone.0015894.

35. Tabar, M.S., Giardina, C., Feng, Y., Francis, H., Sani, H.M., Low, J.K.K., Mackay, J.P., Bailey, C.G., and Rasko, J.E.J. (2022). Unique protein interaction networks define the chromatin remodelling module of the NuRD complex. Febs J 289, 199–214. 10.1111/febs.16112.

36. Abramson, J., Adler, J., Dunger, J., Evans, R., Green, T., Pritzel, A., Ronneberger, O., Willmore, L., Ballard, A.J., Bambrick, J., et al. (2024). Accurate structure prediction of biomolecular interactions with AlphaFold 3. Nature 630, 493–500. 10.1038/s41586-024-07487-w.

37. Schwaiger, M., Mohn, F., Bühler, M., and Kaaij, L.J.T. (2024). guidedNOMe-seq quantifies chromatin states at single allele resolution for hundreds of custom regions in parallel. BMC Genom. 25, 732. 10.1186/s12864-024-10625-3.

38. Parthasarthy, A., and Gopinathan, K.P. (2006). Transcriptional activation of a moderately expressed tRNA gene by a positioned nucleosome. Biochem. J. 396, 439–447. 10.1042/bj20052029.

39. Morse, R.H. (1989). Nucleosomes inhibit both transcriptional initiation and elongation by RNA polymerase III in vitro. EMBO J. 8, 2343–2351. 10.1002/j.1460-2075.1989.tb08362.x.

40. Burnol, A.-F., Margottin, F., Huet, J., Almouzni, G., Prioleau, M.-N., Méchali, M., and Sentenac, A. (1993). TFIIIC relieves repression of U6 snRNA transcription by chromatin. Nature 362, 475–477. 10.1038/362475a0.

41. Li, M., Hada, A., Sen, P., Olufemi, L., Hall, M.A., Smith, B.Y., Forth, S., McKnight, J.N., Patel, A., Bowman, G.D., et al. (2015). Dynamic regulation of transcription factors by nucleosome remodeling. eLife 4, e06249. 10.7554/elife.06249.

42. Perez-Riverol, Y., Bai, J., Bandla, C., García-Seisdedos, D., Hewapathirana, S., Kamatchinathan, S., Kundu, D.J., Prakash, A., Frericks-Zipper, A., Eisenacher, M., et al. (2021). The PRIDE database resources in 2022: a hub for mass spectrometry-based proteomics evidences. Nucleic Acids Res. 50, D543–D552. 10.1093/nar/gkab1038.

43. Ostapcuk, V., Mohn, F., Carl, S.H., Basters, A., Hess, D., Iesmantavicius, V., Lampersberger, L., Flemr, M., Pandey, A., Thomä, N.H., et al. (2018). Activity-dependent neuroprotective protein recruits HP1 and CHD4 to control lineage-specifying genes. Nature 557, 739–743. 10.1038/s41586-018-0153-8.

44. Flemr, M., and Bühler, M. (2015). Single-Step Generation of Conditional Knockout Mouse Embryonic Stem Cells. Cell Reports 12, 709–716. 10.1016/j.celrep.2015.06.051.

45. Álvarez-Salmoral, D., Borza, R., Xie, R., Joosten, R.P., Hekkelman, M.L., and Perrakis, A. (2024). AlphaBridge: tools for the analysis of predicted macromolecular complexes. bioRxiv, 2024.10.23.619601. 10.1101/2024.10.23.619601.

46. Goddard, T.D., Huang, C.C., Meng, E.C., Pettersen, E.F., Couch, G.S., Morris, J.H., and Ferrin, T.E. (2018). UCSF ChimeraX: Meeting modern challenges in visualization and analysis. Protein Sci. 27, 14–25. 10.1002/pro.3235.

47. Dobin, A., Davis, C.A., Schlesinger, F., Drenkow, J., Zaleski, C., Jha, S., Batut, P., Chaisson, M., and Gingeras, T.R. (2013). STAR: ultrafast universal RNA-seq aligner. Bioinformatics 29, 15–21. 10.1093/bioinformatics/bts635.

48. Teissandier, A., Servant, N., Barillot, E., and Bourc’his, D. (2019). Tools and best practices for retrotransposon analysis using high-throughput sequencing data. Mob. DNA 10, 52. 10.1186/s13100-019-0192-1.

49. Zhang, Y., Liu, T., Meyer, C.A., Eeckhoute, J., Johnson, D.S., Bernstein, B.E., Nusbaum, C., Myers, R.M., Brown, M., Li, W., et al. (2008). Model-based Analysis of ChIP-Seq (MACS). Genome Biol. 9, R137. 10.1186/gb-2008-9-9-r137.

50. Liao, Y., Smyth, G.K., and Shi, W. (2014). featureCounts: an efficient general purpose program for assigning sequence reads to genomic features. Bioinformatics 30, 923–930. 10.1093/bioinformatics/btt656.

51. Robinson, J.T., Thorvaldsdóttir, H., Winckler, W., Guttman, M., Lander, E.S., Getz, G., and Mesirov, J.P. (2011). Integrative genomics viewer. Nat. Biotechnol. 29, 24–26. 10.1038/nbt.1754.

52. Love, M.I., Huber, W., and Anders, S. (2014). Moderated estimation of fold change and dispersion for RNA-seq data with DESeq2. Genome Biol. 15, 550. 10.1186/s13059-014-0550-8.

53. Cox, J., Neuhauser, N., Michalski, A., Scheltema, R.A., Olsen, J.V., and Mann, M. (2011). Andromeda: A Peptide Search Engine Integrated into the MaxQuant Environment. J. Proteome Res. 10, 1794–1805. 10.1021/pr101065j.

54. Yu, F., Teo, G.C., Kong, A.T., Fröhlich, K., Li, G.X., Demichev, V., and Nesvizhskii, A.I. (2023). Analysis of DIA proteomics data using MSFragger-DIA and FragPipe computational platform. Nat. Commun. 14, 4154. 10.1038/s41467-023-39869-5.

55. Soneson, C., Iesmantavicius, V., Hess, D., Stadler, M.B., and Seebacher, J. (2023). einprot: flexible, easy-to-use, reproducible workflows for statistical analysis of quantitative proteomics data. bioRxiv, 2023.07.27.550821. 10.1101/2023.07.27.550821.

